# *Ruminococcus bromii* enables the growth of proximal *Bacteroides thetaiotaomicron* by releasing glucose during starch degradation

**DOI:** 10.1101/2022.02.07.479236

**Authors:** Aathmaja Anandhi Rangarajan, Hannah E. Chia, Christopher A. Azaldegui, Monica H. Olszewski, Nicole M. Koropatkin, Julie S. Biteen

**Author notes:** Address correspondence to Julie S. Biteen.

## Abstract

Complex carbohydrates shape the gut microbiota and the collective fermentation of resistant starch by gut microbes positively affects human health through enhanced butyrate production. The keystone species *Ruminococcus bromii* (*Rb*) is a specialist in degrading resistant starch; its degradation products are used by other bacteria including *Bacteroides thetaiotaomicron* (*Bt*). We analyzed the metabolic and spatial relationships between *Rb* and *Bt* during potato starch degradation and found that *Bt* utilizes glucose that is released from *Rb* upon degradation of resistant potato starch and soluble potato amylopectin. Additionally, we found that *Rb* produces a halo of glucose around it when grown on solid media containing potato amylopectin and that *Bt* cells deficient for growth on potato amylopectin (Δ*sus Bt*) can grow within the halo. Furthermore, when these Δ*sus Bt* cells grow within this glucose halo, they have an elongated cell morphology. This long-cell phenotype depends on the glucose concentration in the solid media: longer *Bt* cells are formed at higher glucose concentrations. Together, our results indicate that starch degradation by *Rb* cross-feeds other bacteria in the surrounding region by releasing glucose. Our results also elucidate the adaptive morphology of *Bt* cells under different nutrient and physiological conditions.

**Impact Statement:** Dietary intake of complex carbohydrates including resistant starch benefits human health by supporting the growth of keystone species that increase the microbiome diversity via cross-feeding. For instance, *Ruminococcus bromii* (*Rb*) is a specialist in degrading resistant starch, and the byproducts of this degradation process are used by bacterial species. In this study, we show that *Bacteroides thetaiotaomicron* (*Bt*) cross-feeds on the glucose released during potato starch degradation by *Rb*. We also show that proximity to *Rb* cells that are releasing sugars is important for the growth of the cross-fed *Bt* in solid media. Additionally, we find a longer phenotype for *Bt* cells grown on solid media in high glucose conditions, indicating that, like other bacteria, human gut bacteria undergo significant nutrient-dependent morphological adaptations.

## Introduction

The human gut microbiome consists of trillions of microbes, most belonging to the Firmicutes and Bacteroidetes phyla (1,2). The health benefits provided to the host by a well-balanced gut microbiota include digestion of complex polysaccharides, synthesis of micronutrients, resistance against pathogens, and development of immunity (3,4). The microbial composition varies across different regions of the gut and this composition is highly organized into distinct biogeographies depending on nutrient availability (5–9). The formation of these coexisting bacterial communities results from an intricate synergy between members of the gut microbiome during carbohydrate breakdown (10–12). For example, mucin cross-feeding has been reported between infant *Bifidobacteria* and *Eubacterium halii* (13). *Bifidobacterium pseudolongum* degrades Hi-Maize resistant starch whose byproducts are used by other bifidobacterial species (14). Cross feeding has also been observed in *Bifidobacterium longum, Anaerostipes caccae*, and *Roseburia intestinalis* during growth on oligofructose (15). However, there exists a fundamental gap in our knowledge about the metabolic and spatial relationships between gut microbiome members during cross-feeding at the single-cell level.

Diet is a major factor that influences the gut microbiota, and it can cause short- and long-term changes in gut microbial communities (16–18). Starch is the most abundant polysaccharide in the Western diet (19). Resistant starch is indigestible by humans but can be utilized by the microbiota; this metabolism creates short chain fatty acids like butyrate, which confer enormous health benefits to the host (20–24). The Firmicutes *Ruminococcus bromii* (*Rb*) has evolved a unique extracellular multi-protein enzymatic machinery known as the amylosome. Akin to cellulosome complexes synthesized by cellulose-degrading bacteria and fungi, amylosomes are comprised of dockerin-containing enzymes that bind cohesin domains on scaffoldin proteins, and this protein complex efficiently binds and degrades resistant starch (25–28). *Rb* is one of the few gut bacteria that can degrade resistant starch thus it acts as a “keystone” species that supports other species by releasing sugars and other metabolites (25,26,29,30). *Bacteroides thetaiotaomicron (Bt)* is another prominent member of the gut microbiome, but unlike *Rb, Bt* cannot break down resistant starch. However, *Bt* does have the ability to catabolize several types of glycans, including soluble polysaccharides via the expression of discrete, polysaccharide utilization loci (PULs) of which *sus* (the starch utilization system) is the best studied (31–33). Though *Bt* can utilize a wide variety of dietary or host-derived glycans, the availability of a specific carbon source to *Bt* is shaped by the local gut environment (6,34). When grown in co-culture with *Rb, Bt* can cross-feed on the resistant corn starch degradation products of *Rb*. However, the metabolic profile of the sugars that *Rb* provides to *Bt* during potato starch degradation is unknown. As *Rb* and *Bt* reside in the gut environment, it is also imperative to understand how the distance between *Rb* and *Bt* on a solid surface affects cross-feeding.

The spatial properties of a microbial community also depend strongly on the environment. Bacteria possess unique cell morphologies that help them survive in their native environments (35). For instance, large bacteria are typically found in nutrient-rich environments such as insect guts and marine sediments(36). Several bacterial species, including gut bacteria, change their size and shape to adapt to various nutrient and environmental conditions (37–39). These pleomorphic bacteria regulate their cellular machinery to form longer phenotypes to survive in response to a physiological or environmental change. Uropathogenic *Escherichia coli, Legionella pneumophila*, and *Streptococcus pneumoniae* form long filaments to evade the host immune response and enhance attachment to host cell surfaces(40–42). Other bacteria such as *Proteus mirabilis* and *Vibrio parahaemolyticus* sense solid surfaces through flagella and elongate to promote swarming motility (43,44). How gut anaerobes adapt to different nutrient and environmental conditions at the single-cell and community level is not well-studied (39).

The ability of *Rb* to degrade resistant corn starch and soluble starch and cross-feed other bacteria including *Bt* has been described (25,29). However, we lack a detailed understanding of the metabolic and spatial relationships between *Rb* and other bacteria that support this cross-feeding. In this study, we show that *Rb* releases glucose as the major byproduct of both resistant potato starch and soluble potato amylopectin degradation. Furthermore, we show that *Bt* growth on solid media is enhanced near *Rb* that is releasing the byproducts of potato amylopectin degradation. Moreover, with microscopy we show that cross-fed *Bt* has a longer phenotype in proximity to *Rb* when compared to *Bt* that is distant from *Rb* due to the local prevalence of released glucose. This model system we have constructed on solid media provides a lens into bacterial cross-feeding in the gut environment.

## Methods

### Bacterial strains and growth conditions

*Δsus UnaG Bt* was generated by counter-selectable allelic exchange in a *ΔsusA-G Bt* (*Δtdk*) thymidine kinase deletion mutant with codon optimized UnaG (45) and grown as previously described (46). *Bt* ATCC 29148 (VPI-5482) and its derivative *Δsus UnaG Bt* were grown in TYG medium (47) or minimal medium (48). Bilirubin was added in the media to final concentration of 25 μM. Carbon sources were added to 5 mg/ml unless otherwise stated. *Rb* L2-63 was grown in modified RUM medium (26), consisting of (per 100 ml) tryptone (0.5 g), yeast extract (0.25 g), resazurin (50 μg), hematin (30 μM), and L-histidine (3 mM), salt mixture consisting of NaHCO_3_ (0.4 g), L-cysteine (0.1 g), (NH_4_)_2_SO_4_ (0.09 g), K_2_HPO_4_ (0.045 g), KH_2_PO_4_ (0.045 g), NaCl (0.09 g), MgSO_4_ (0.004 g), and CaCl_2_ (0.009 g), and a vitamin mixture consisting of biotin (20 μg), cobalamin (20 μg), p-aminobenzoic acid (60 μg), folic acid (100 μg), pyridoxamine (300 μg), thiamine (100 μg), riboflavin (100 μg), D-pantothenoic acid hemicalcium salt (100 μg), and nicotinamide (100 μg). A short chain fatty acid (SCFA) mixture consisting of acetic acid (63.7 mM), propionic acid (17.8 mM), isobutyric acid (5.75 mM), isovaleric acid (1.95 mM), and valeric acid (1.95 mM) was also added. The pH was set to 7.4 ± 0.2 using 6 M NaOH. The medium was filter sterilized and reduced in the anaerobic chamber before inoculation. 2× RUM medium was diluted with an equal volume of a carbohydrate solution containing maltose (final concentration 5 mg/ml), fructose (final concentration 5 mg/ml) and glycogen (final concentration 2.5 mg/ml) or potato amylopectin (final concentration 5 mg/ml) to grow the cells. Resistant potato starch (Bob’s Red Mill^®^ raw potato starch, unmodified) was sterilized with 70% ethanol twice and air dried before use (final concentration 25 mg/ml). All strains were grown at 37 °C under anaerobic conditions (5% H_2_, 85% N_2_, 10% CO_2_) in a anaerobic chamber. The OD_600_ was measured using a spectrophotometer (Genesys 20, Thermo Fisher Scientific) or absorbance reader (PowerWave HT, Biotek Instruments). Data were recorded using Gen5 software (BioTek Instruments) and Prism (GraphPad).

### Growth of bacteria in solid media

A modified RUM media was used for growing *Rb* and *Bt* in solid media. For 100 ml media, 95 ml of water was mixed with tryptone (0.5 g), agarose (2 g) and the salt mixture as described in the previous section except NaHCO_3_ and autoclaved. After autoclaving this media, a filter sterilized solution consisting of NaHCO_3_, L-cysteine, resazurin, hematin, and L-histidine, the vitamin mixture, and the SCFA mixture were added in the final concentration as described in the previous section. The pH was set to 8 ± 0.2 using 6 M NaOH. Potato amylopectin was used at 5 mg/ml for growing *Rb*. For growing *Bt*, glucose, ribose, xylose, or maltose was used at 200 μM, 2 mM, or 20 mM final concentration, respectively. Potato amylopectin was used at final concentration of 0.4, 2, or 10 mg/ml. The plates were reduced for 48 hrs inside the anaerobic chamber. *Rb* cells were washed with modified RUM media without carbon source and *Bt* cells were washed with minimal media without carbon source before plating. 10 μl of *Rb* and *Bt* cells were plated and incubated for 72 hrs before taking the plate pictures and determining glucose concentration. After 24 hrs incubation, in plates previously inoculated with *Rb* at the center, *Bt* was plated at different distances from *Rb* (0.5, 1, 1.5, and 2 cm) and incubated for 72 hrs before taking plate pictures (Color QCount, Spiral Biotech).

### Determination of glucose concentration in agarose plates

For the determination of glucose in the RUM media plates, 1 cm^2^ samples were cut from the agar, dissolved in 2× volume 6 M sodium iodide and heated at 55 °C for 15 min. The dissolved agarose solution was used to determine the concentration of glucose according to manufacturer’s instruction (D-Glucose assay kit (GOPOD format, Megazyme).

### Isolation of Spent RUM media

*Rb* was inoculated in modified RUM media with resistant potato starch (final concentration 25 mg/ml) or potato amylopectin (final concentration 5 mg/ml) as a carbon source. After 72 hrs of growth, the samples were centrifuged at 5000 rpm for 5 minutes. The supernatant spent RUM was removed and filter sterilized with 0.2 μm filter. To this spent RUM, 1× RUM salt mixture solution mentioned above was added to achieve pH 7 before inoculating with *Bt*.

### Analysis of sugars using high-performance anion exchange chromatography with pulsed amperometric detection (HPAEC-PAD)

HPAEC-PAD analysis of soluble sugars was done using ICS-6000 system (Thermo Scientific Dionex). Samples were separated with a CarboPac PA-100 anion exchange column (250 mm × 2 mm; Thermo Fisher Scientific) and a CarboPac PA-100 guard column (Thermo Fisher Scientific). Detection was enabled by PAD with a gold working electrode and an Ag/AgCl reference electrode using a standard AAA potential waveform. Each sample was run on the column for 40 min at a constant flow rate of 1 ml/min. A gradient composed of the following eluents was used at 25 °C: (buffer A) 0.1 M sodium hydroxide, (buffer B) 0.1 M sodium hydroxide + 0.5 M sodium acetate. Before every run, the column was washed with 100% buffer A for 10 min. For monosaccharide and oligosaccharides, the following gradient was applied after injection: 0 – 5 min, isocratic 100% buffer A; 5 – 24 min linear gradient to 50% buffer A and 50% buffer B; 24 – 34 min 100% buffer B. For polysaccharides, the following gradient was applied after injection: 0 – 60 min, linear gradient to 90% buffer B and 10% A; 61 – 74 min 90% buffer A and 10% buffer B. 1 mM of glucose, maltose, maltotriose, maltotetraose, and 5 mg/ml potato amylopectin were used as standards. 3, 1.5, 0.75, 0.375, and 0.187 mM glucose standards were used for the standard calibration curves. Data integration and analysis were performed using the Chromeleon 7.2 chromatography data system (Thermo Scientific Dionex).

### Sample preparation and live-cell imaging

Samples were placed on gridded coverslip (28 mM, 50 μm grids, IBIDI) and sealed with epoxy (5 Minute Epoxy®, Devcon) to maintain cells in anaerobic condition and to enable live-cell imaging (49). Phase-contrast images were taken on an Olympus IX71 inverted fluorescence microscope using a 1.40 numerical aperture 100× wide field oil-immersion objective. *UnaG Bt* and *Δsus UnaG Bt* cell samples were imaged using a 488-nm laser (Coherent Sapphire 488–50; 8 – 18 W/cm^2^) with 488-nm long pass filter, and fluorescence emission was detected on a 512 × 512 pixel Photometrics Evolve electron-multiplying charge-coupled device camera at 25 frames/s.

### Cell length measurements

Cell segmentation and quantification of cell dimensions were done with a custom Python script. Phase masks were produced using an in-house trained cell segmentation model implemented with Cellpose (50). Erroneous segmentations were manually corrected using the Cellpose GUI. Cell lengths and widths were determined from the long- and short-axis of each cell in the phase mask using the scikit-image Python package (51). Mean lengths and widths were calculated and *P*-values were determined using one-way ANOVA and post-hoc Tukey tests with 95% confidence intervals.

## Results

### The major byproduct of resistant potato starch degradation by Rb is glucose, which is then utilized by Bt

We investigated the ability of *Bt* to cross-feed on the byproducts of resistant potato starch degradation by *Rb* based on previous studies that showed that *Bt* can grow on the products of resistant corn degradation by *Rb* in co-culture (25). While *Rb* can grow on starch and malto-oligosaccharides, most strains including the *Rb* L2-63 used in our study cannot grow on glucose (27). We isolated the spent medium after *Rb* growth on resistant potato starch by filtering out the *Rb* cells. Growth of *Rb* on resistant starch acidifies the medium, which was initially at pH 7.4±0.2, to pH 5 after 72 hrs of growth (Figure S1a). Despite the presence of cross-feeding sugars from *Rb*, the acidic pH inhibits *Bt* growth consistent with previous findings (Figure S1b)(52). However, after it is neutralized to pH 7, *Bt* can grow in the spent medium of *Rb* with no additional carbohydrate source due to the presence of the sugars released by *Rb* (Figure 1a). We identified the sugars released by *Rb* with high-performance anion exchange chromatography with pulsed amperometric detection (HPAEC-PAD), which showed that glucose is the major byproduct of the resistant starch degradation of potato starch by *Rb*. This glucose is utilized by *Bt* during growth in the neutralized spent medium after 24 hrs of growth (Figure 1b). Next, we quantified the amount of glucose released as *Rb* grows in resistant potato starch. The glucose concentration increases with time as *Rb* degrades resistant potato starch and reaches ∼1 mM after 96 hrs (Figures 1c and S2a). No maltose, malto-oligosaccharides, or soluble polysaccharides were identified in our analysis; if generated, these may be rapidly utilized by *Rb* instead of glucose (Figure S2b and S2c). These results show that *Rb* releases glucose as the major byproduct of resistant potato starch degradation, that the amount of glucose in the medium increases over time, and that this glucose is utilized by *Bt*.

**Figure 1.**
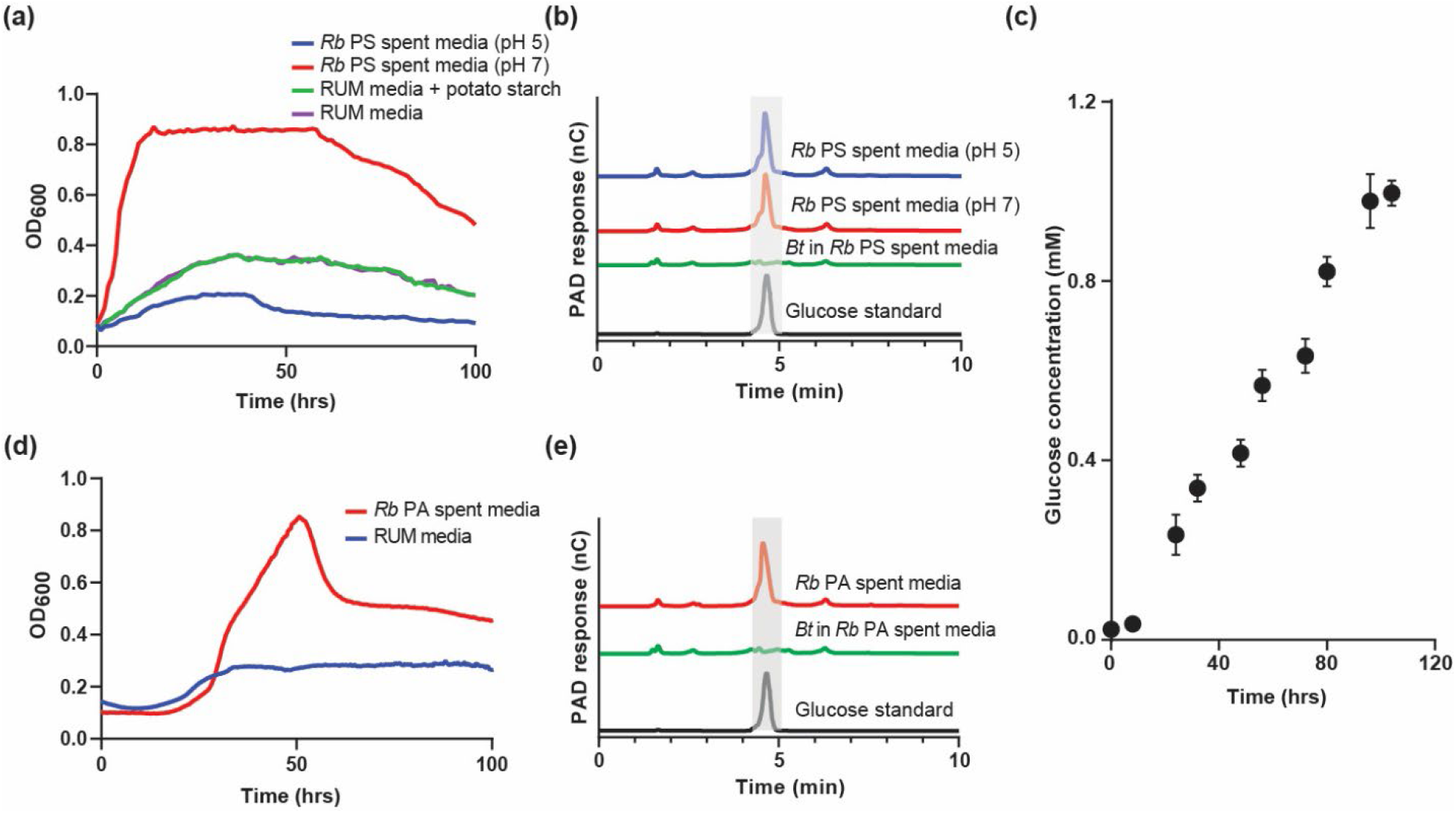
(a) Growth of *Bt* cells in *Rb* resistant potato starch (PS) spent medium (pH 5, blue), neutralized *Rb* potato starch spent medium (pH 7, red), RUM medium with resistant potato starch (green), and RUM medium alone (magenta) measured by absorbance at 600 nm. Mean values of three biological replicates are shown. (b) Analysis of the spent medium by HPAEC-PAD. Shaded grey region on the traces highlight where the glucose peak should be if glucose is present. Black, blue, red and green traces represent the glucose standard, *Rb* resistant potato starch spent medium at pH5, *Rb* resistant potato starch spent medium at pH7 and *Rb* resistant potato starch spent medium at pH7 after the growth of *Bt* cells, respectively. (c) Glucose concentration in *Rb* resistant potato starch spent medium isolated at different time points (0 – 104 hrs). Three biological replicates were used for this HPAEC-PAD analysis. Error bars indicate standard error of the mean. (d) Growth of *Δsus unaG Bt* cells in neutralized *Rb* spent medium isolated from *Rb* cells grown in potato amylopectin (PA, red) and the RUM medium control (blue) measured by absorbance at 600 nm. (e) Analysis by HPAEC-PAD of the *Rb* spent medium isolated from *Rb* cells grown in potato amylopectin. Shaded grey region on the traces highlight where the glucose peak should be if glucose is present. Black, red and green traces represent the glucose standard, *Rb* potato amylopectin spent medium and *Rb* potato amylopectin spent medium after the growth of *Δsus unaG Bt* cells, respectively.

### Spatial proximity to the glucose released by Rb on potato amylopectin agarose helps Δsus UnaG Bt growth

Gut bacteria are confined to spatially distinct regions of the intestine and can be tethered to the solid intestinal surface (6,8). To study the cross-feeding between *Rb* and *Bt* in a spatial context, we used solid agarose plates with potato amylopectin as a carbon source, since the cloudiness of the insoluble resistant starch makes cells grown on this source unsuitable for microscopy analysis. *Rb* grows on potato amylopectin similar to growth on fructose and glycogen as carbon source (Figure S3a). Since *Bt* can also grow on amylopectin, we used a *Δsus Bt* strain in which the *sus* operon (*susA-G*) is deleted, and tagged it with the anaerobic fluorescent protein UnaG, which fluoresces in the presence of a bilirubin cofactor when excited at 488 nm (45). The constructed *Δsus UnaG Bt* strain cannot grow on potato amylopectin but can grow on glucose and fluoresces in the presence of a bilirubin cofactor when excited at 488 nm (Figure S3b and c). This *Δsus UnaG Bt* strain can also grow on the neutralized spent medium of *Rb* grown in potato amylopectin (Figure 1d). HPAEC-PAD detected that *Rb* releases glucose after utilizing potato amylopectin similar to resistant potato starch, and that this glucose is utilized by *Bt* (Figure 1e). Like in liquid cultures, we confirmed that *Rb* can grow on agarose plates supplemented with amylopectin whereas *Δsus UnaG Bt* cannot (Figure S4). Moreover, *Rb* growing on potato amylopectin plates form a ‘halo’ of glucose (Figure 2a). The halo diameter and the total amount of glucose detected decrease with *Rb* dilution (Figure 2b). However, the amount of glucose per square centimeter remains constant across different dilutions of *Rb* (Figure S5).

**Figure 2.**
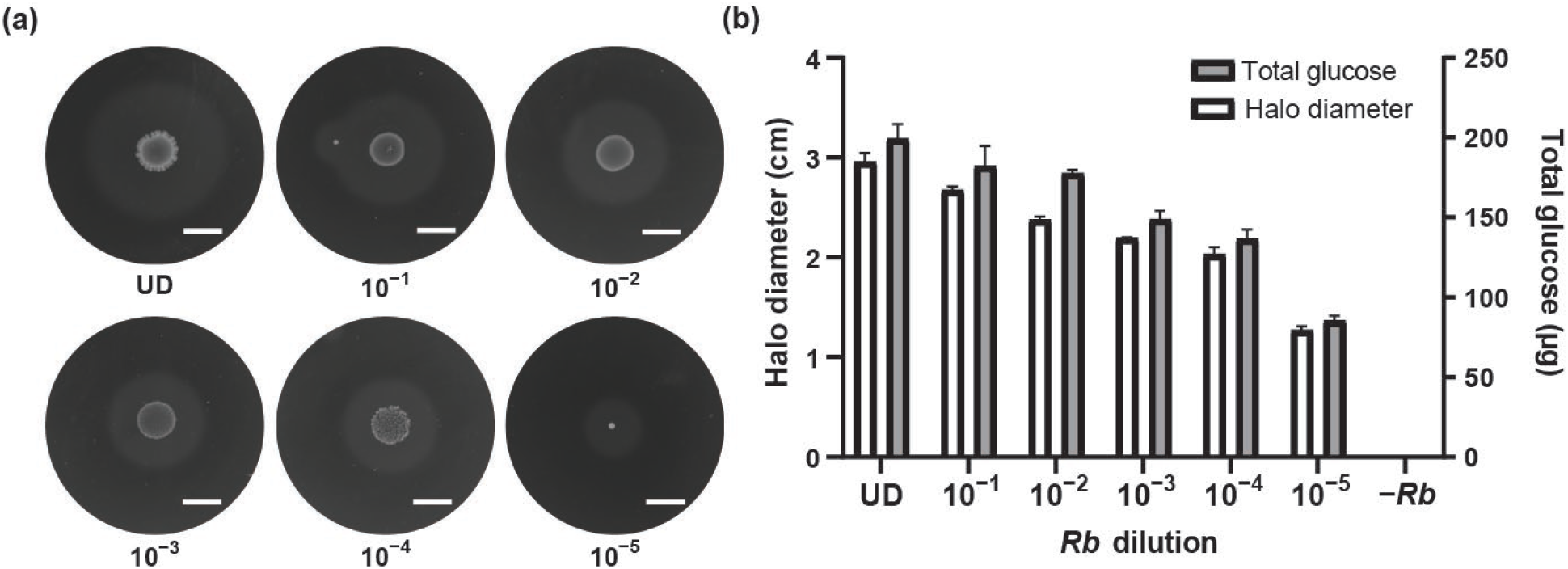
(a) Pictures of *Rb* cells plated on agarose plates at dilutions from undiluted (UD) to 10^−5^ with potato amylopectin (5 mg/ml) showing a halo around the cells. Scale bars: 1 cm. (b) Halo diameter (white bars) and total mass of glucose in the halo (grey bars) for each *Rb* dilution. The average values of two biological replicates are shown. Error bars indicate standard error of the mean.

Next, we spotted *Δsus UnaG Bt* at different distances (0.5 – 2 cm) from *Rb* to study the effect of their spatial proximity to *Rb*. Across different dilutions, we measured *Δsus UnaG Bt* growth from the absorbance intensity, which is proportional to the colony density. *Δsus UnaG Bt* grew better within the halo of glucose released by *Rb* relative to *Δsus UnaG Bt* outside the halo (Figure 3a-c). The percentage area of the *Bt* colony inside the halo linearly correlates with the intensity of the colony observed across all dilutions (Figure 3d). These results show that spatial proximity to *Rb* releasing glucose results in better growth of *Δsus UnaG Bt* due the local availability of glucose released by *Rb*.

**Figure 3.**
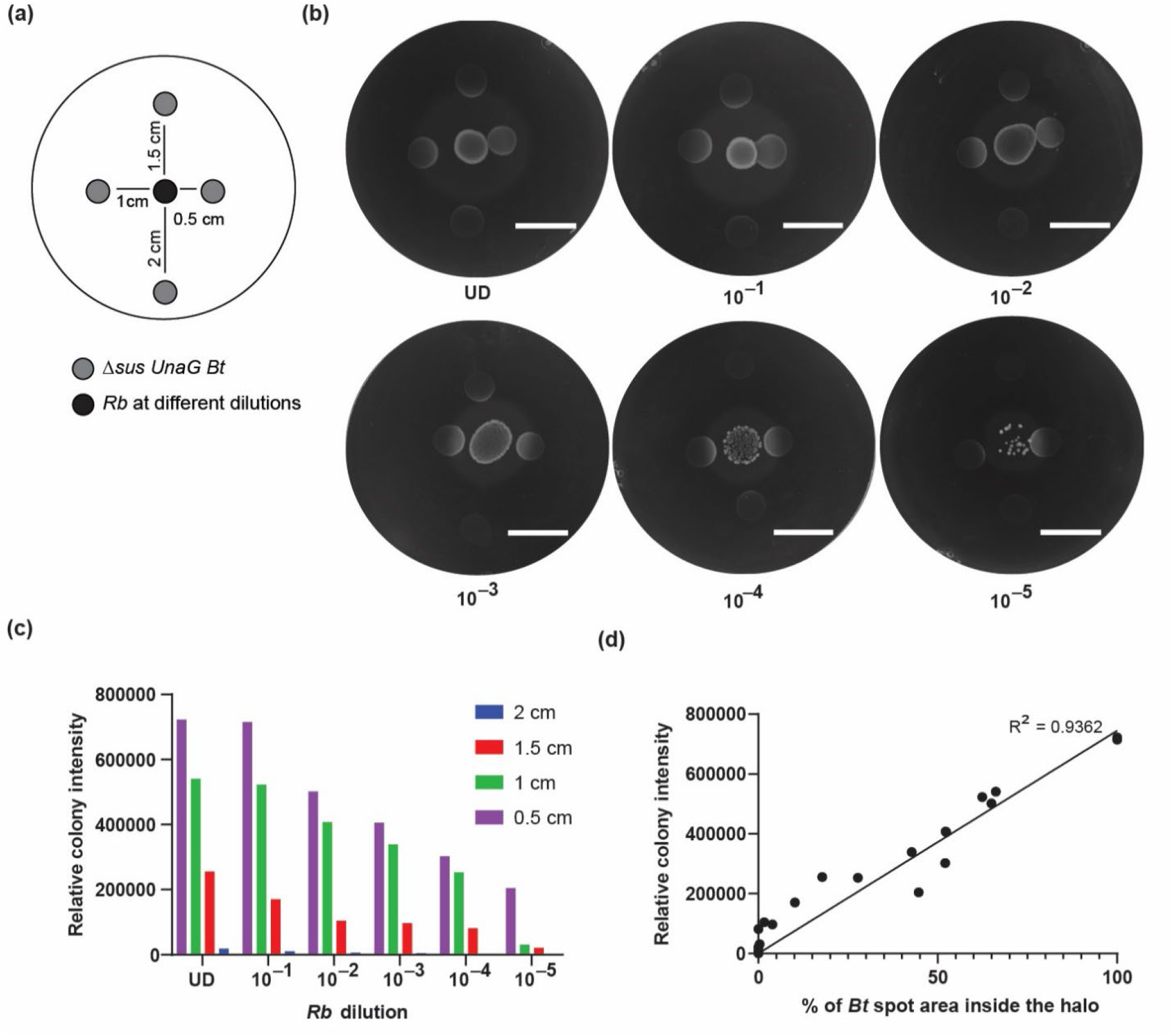
(a) Schematic of the potato amylopectin plates prepared to analyze the spatial relationship between *Δsus UnaG Bt* and *Rb*: 10 μl *Δsus UnaG Bt* was spotted at four different distances from a central spot of *Rb*. (b) Pictures of cells plated on agarose plates with potato amylopectin (5 mg/ml). The central *Rb* cells are plated at dilutions from undiluted (UD) to 10^−5^ and a grey halo is observed around this spot. *Δsus UnaG Bt* are cells spotted at 0.5, 1, 1.5, and 2 cm from the *Rb*. Scale bars: 2 cm. (c) Growth of each *Bt* spot measured as the intensity of the spot relative to the background intensity. (d) We calculated the percentage area inside the halo for each *Δsus UnaG Bt* spot and compared it to the normalized intensity of that *Δsus UnaG Bt* spot. Data points corresponding to all *Rb* dilutions and all distances of *Δsus UnaG Bt* are represented as dots. Black line: fit to a linear curve.

### Single-cell analysis reveals a longer Δsus UnaG Bt phenotype near Rb releasing glucose on solid media

To analyze the cell morphology of *Δsus UnaG Bt* cells inside and outside the halo, we imaged the cells at 0 and 6 hrs using the gridded coverslip as reference for unbiased sampling. We previously reported an elongated phenotype (2.3 μm) for *Bt* cells grown in liquid medium under sugar-limited conditions in the presence of sodium carbonate (39). Hence, we hypothesized that *Δsus UnaG Bt* cells grown within the glucose halo of *Rb* would have the wild-type phenotype, while *Δsus UnaG Bt* in the sugar-limited region outside the halo would have the elongated phenotype. However, contrary to our expectations, *Δsus UnaG Bt* grown inside the halo containing glucose has a longerphenotype (2.4 μm) compared to the cells outside the halo (1.9 μm) after 6 hrs of incubation (Figures 4a-b and S6a). The longer cells inside the halo are similar to the *Δsus UnaG Bt* cells plated on glucose (125 μM) in terms of size and density (Figures 4a-b and S6b). On the other hand, the length of *Δsus UnaG Bt* cells grown in liquid medium with glucose (27 mM), the concentration typically used for growing *Bt*, showed no difference in the length of the cells after 6 hrs of incubation (Figure S7). These results show that *Δsus UnaG Bt* cells on solid media have a longer morphology in the vicinity of *Rb* releasing glucose.

**Figure 4.**
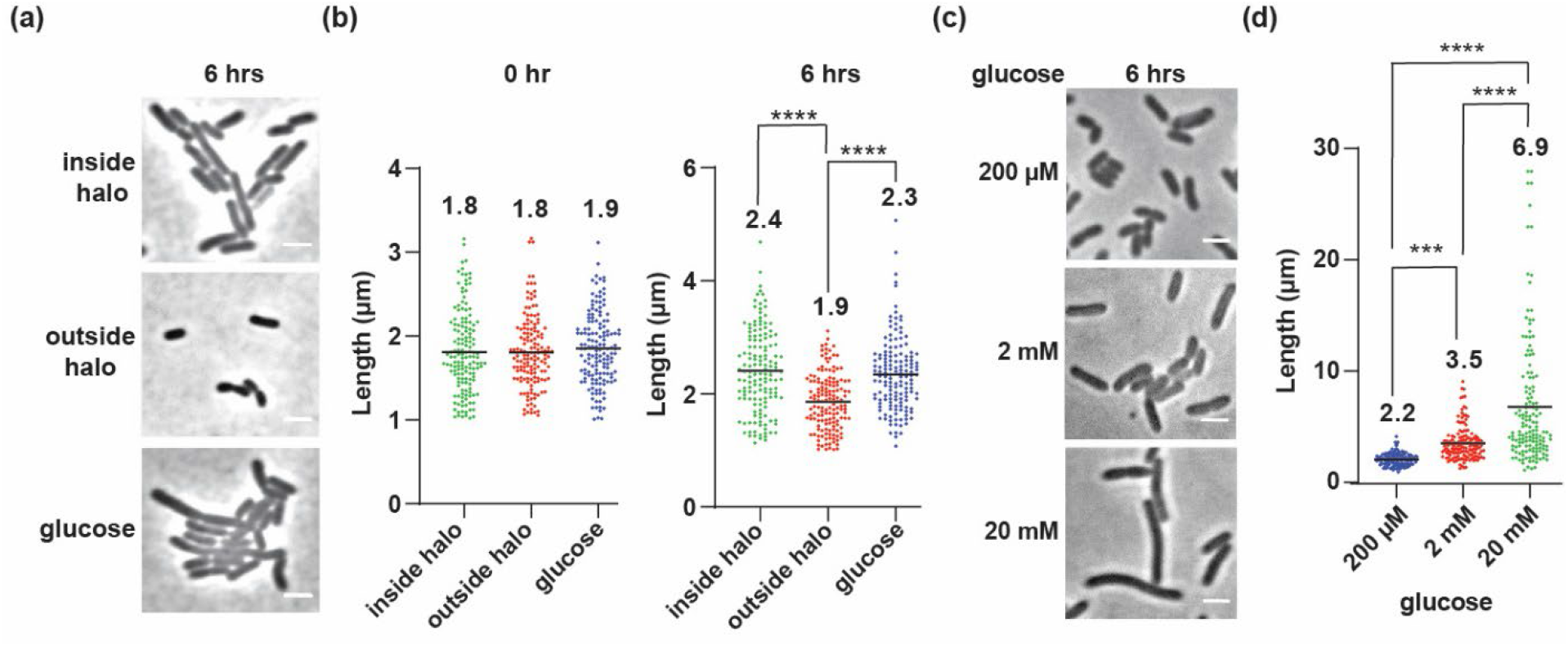
(a) Representative phase-contrast microscopy images of *Δsus UnaG Bt* cells inside the glucose halo, outside the glucose halo, and on plates with glucose at 6 hrs. Scale bars: 2 μm. (b) Length of *Δsus UnaG Bt* cells grown in inside the glucose halo (green), outside the glucose halo (red), and on plates with glucose (blue) at 0 and 6 hrs. Each circle represents the length of a single cell from a set of three biological replicates (*n* = 150 cells). Mean value is indicated by black line. Statistical significance was determined by one-way ANOVA and post-hoc Tukey tests. Significant differences are indicated by **** (*P* < 0.0001) (c) Representative phase-contrast microscopy images of wild-type *Bt* cells taken after 6 hrs incubation on plates with 200 μM, 2 mM, and 20 mM glucose. Scale bars: 2 μm. (d) Length of wild-type *Bt* cells grown on 200 μM (blue), 2 mM (red) and 20 mM (green) glucose for 6 hrs. Each circle represents the length of a single cell from a set of three biological replicates (*n* = 150 cells). Statistical significance was determined by one-way ANOVA and post-hoc Tukey tests. Significant differences are indicated by **** (*P* < 0.0001) and *** (*P* < 0.001).

### The longer Bt morphology in solid media depends on the glucose concentration

To test whether this longer morphology is specific to glucose alone or a general phenomenon, we analyzed the morphology of wild-type *Bt* cells grown on solid media with other monosaccharides (xylose or ribose), the disaccharide maltose, or the polysaccharide potato amylopectin. *Bt* uses unique set of genes to transport and metabolize each of these sugars (34,53–55). Wild-type *Bt* cells was plated on solid media with different concentrations of sugar (200 μM, 2 mM, or 20 mM) and on potato amylopectin (0.4, 2, or 10 mg/ml) and imaged after 6 hrs of incubation. In glucose, the cell length was 1.8 μm at the time of incubation (0 hr), which increased to 2.2 μm at 200 μM after 6 hrs of incubation (Figure 4c-d). The average length was yet higher (3.5 μm) after incubation in 2 mM glucose, and incubation at 20 mM produced very long cells (average length of 6.9 μm with some cells as long as 27 μm) (Figures 4c-d and S8). In all the mono- and disaccharides tested, the cells at 20 mM sugar concentration were significantly elongated relative to cells at 200 μM sugar concentration (Figure S9a-c). A similar increase in cell size was seen for cells grown at 10 mg/ml amylopectin (2.2 μm) compared to 0.4 mg/ml (1.9 μm) (Figure S9d). However, contrary to what was observed with glucose, we did not see a consistent increase in cell length as a function of concentration for xylose, ribose and potato amylopectin. At 20 mM, the very long *Bt* cells seen at glucose were seen only for few outliers in maltose and were never seen in similar concentrations of xylose, ribose, or potato amylopectin (Figure S9). These results show that *Bt* adapts its morphology at higher concentrations of different sugars in solid media, but that the concentration-dependent increase in cell size, including very long cells at 20 mM, is seen only in glucose.

## Discussion

*Rb* has evolved to possess specialized proteins that form amylosome complexes that efficiently bind and degrade resistant starches, and *Rb* has been established as a keystone species in the gut microbiota because it produces starch degradation byproducts that cross-feed other community members (25–29). Studying the mechanism of this metabolism and resource sharing will provide important information toward understanding community formation and maintenance. In this study, we show that *Rb* releases glucose upon degradation of resistant starch and soluble potato amylopectin (Figure 1). On a solid surface, the glucose released in the vicinity of *Rb* can be utilized by *Bt* (Figure 3). Our results complement recent findings which showed that *Rb* possesses several surface-anchored and secreted amylosome complex proteins with starch-binding and starch-degrading capacity (26,27). These proteins could take in binding and degrading starch in our system to produce the glucose byproduct that we observe as a halo of glucose around *Rb* (Figures 2a and 3b). We show the total glucose amount and the halo diameter decrease with higher dilutions of *Rb* (Figures 2 and 3b). This decrease in total glucose at higher *Rb* dilutions is likely due to there being fewer amylosome complex present when the *Rb* cell density is lower. This degradation outside the *Rb* cell can help other proximal bacteria, like *Bt*, allowing them to utilize these byproducts and thrive in the complex gut environment. Further studies are needed to elucidate the mechanism and dynamics of resistant starch degradation.

In our study, HPAEC-PAD analysis of the products of resistant potato starch degradation by *Rb* showed that glucose is the only major byproduct; we did not identify any other oligosaccharides or polysaccharides (Figure S2b-c). A previous study of the degradation of Hi-Maize resistant starch by *Rb* by nuclear magnetic resonance (NMR) spectroscopy detected minor amounts of maltose and maltotetraose in addition to glucose (29). This difference may be due to differences between potato starch and Hi-Maize substrates or differences in sensitivity between HPAEC-PAD and NMR. However, the absence of anything but glucose in our assays indicates that *Rb* immediately utilizes any oligomeric byproducts as they are released. Alternatively, since the *Rb* amylosome complex contains several dockerin-containing proteins that are capable of binding longer and shorter oligosaccharides (26–28), it is also plausible that any oligosaccharides byproducts that form from resistant starch degradation are bound by dockerin proteins and filtered out of the spent medium.

Morphological pleomorphism and plasticity has been observed in several bacterial species in response to variety of nutrient and environmental conditions (56). We previously observed elongated *Bt* cells (2.3 μm) in nutrient-poor conditions in liquid medium in the presence of sodium carbonate (39). However, contrary to our expectations, when *Bt* is grown on solid media, we observe an elongated phenotype at higher sugar concentrations; this result is consistent across a variety of sugars: glucose, maltose, xylose, ribose, and potato amylopectin polysaccharide (Figures 4d and S9). This effect is more pronounced in glucose: the *Bt* cell size increases with glucose concentration, and very long *Bt* cells (mean length 6.9 μm) grow at the highest glucose concentration of 20 mM (Figure 4c-d). This result indicates that the *Bt* cell elongation machinery might be regulated by glucose similar to observations in *E. coli* and *B. subtilis* in which the cell elongation machinery is regulated by uridine diphosphate glucose through FtsZ (36,57,58). The solid surface affects the *Bt* cell elongation machinery, similar to other bacteria which form long filamentous structures upon attachment to solid surface (40,41,43,44). We observe a differential regulation of the cell division machinery for *Bt* in solid and liquid media under different nutrient conditions. The underlying molecular mechanisms involved in the differential phenotypic responses to nutrient availability under varied nutrient and environmental conditions require further investigation. The adaptation of *Bt* to various nutrient conditions both in liquid medium and on solid surfaces may allow this bacterium to effectively compete with other bacteria and persist within the gut ecosystem.

## Authors and Contributors

A.A.R., H.E.C., N.M.K. and J.S.B. conceived the study and designed the experiments. A.A.R. conducted all the experiments with assistance from M.H.O. C.A.A. curated and analyzed cell length measurements. A.A.R. and J.S.B. wrote the manuscript. All the authors revised the final version of the manuscript.

## Funding information

This work was supported by Army Research Office grant W911NF-18-1-0339 to J.S.B. and

N.M.K.

## Acknowledgements

We thank Gabriel V. Pereira for his assistance in using HPAEC-PAD. We thank Eric C. Martens for access to BioTek plate reader and Dionex ICS-6000 system. We also thank Supriya Suresh Kumar for the construction of the Δ*susA-G* mutant. We are thankful to the Mobley lab for access to their Color QCount.

