## Supplemental Data 1 for "*Ruminococcus bromii* enables the growth of proximal *Bacteroides thetaiotaomicron* by releasing glucose during starch degradation"

**Contents:**

Supplementary Figures S1 – S9

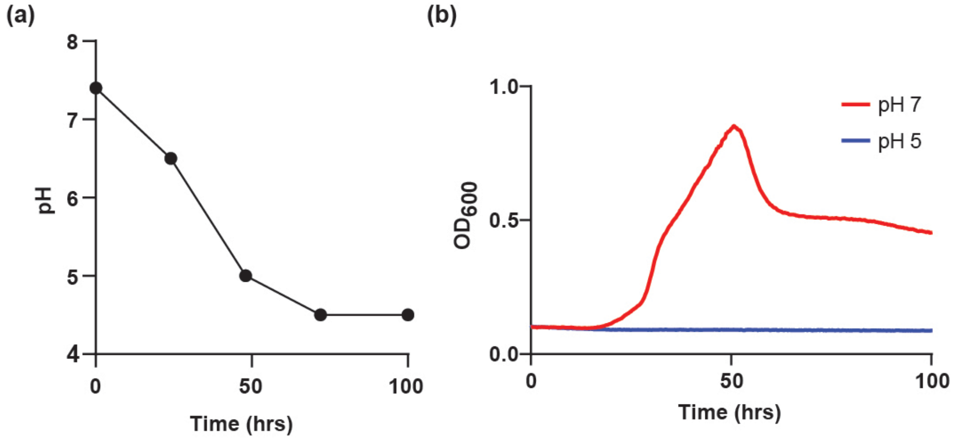


**Figure S1.** (a) Measurements of the pH of the *Rb* resistant potato starch spent medium after 0, 24, 48, 72, and 100 hrs of growth. (b) Growth of *Bt* cells in minimal medium with glucose (5 mg/ml) as a carbon source at pH 7 (red) and pH 5 (blue) measured by absorbance at 600 nm.

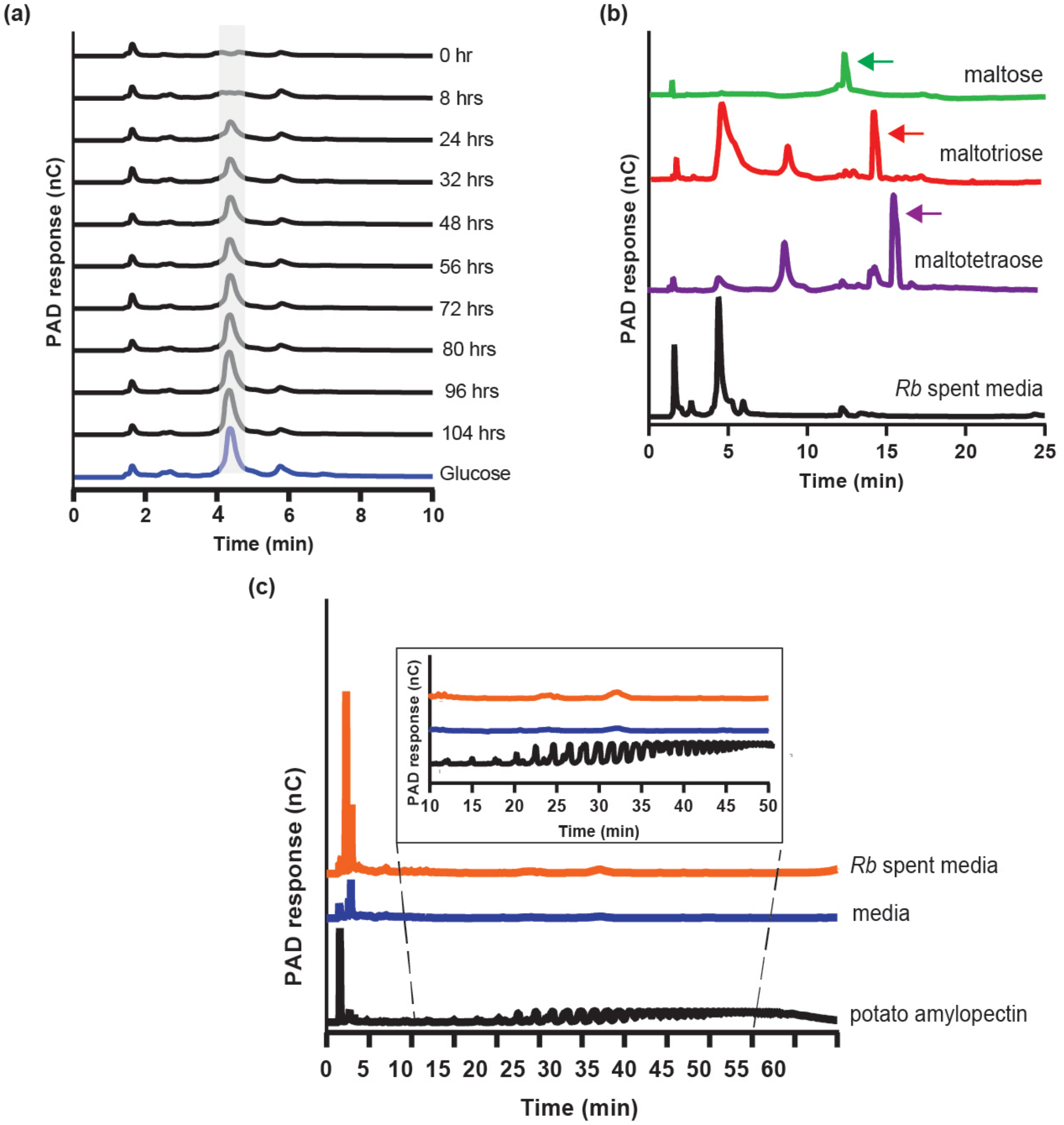


**Figure S2.** (a) Analysis of *Rb* spent medium at different time points (0 – 104 hrs) along with a 1 mM glucose standard using HPAEC-PAD. Traces show a peak (shaded grey region) corresponding to glucose. (b) Analysis of *Rb* spent medium alongside maltose, maltotriose, and maltotetraose standards (1 mM). The green, red, and purple arrows indicate the maltose, maltotriose and maltotetraose peaks, respectively; these peaks are not detected in the spent medium. (c) Analysis of *Rb* spent medium (orange) with potato amylopectin polysaccharide (5 mg/ml; black) and medium control (blue). The inset shows the time points from 10 – 50 min, indicating glucose polymers in the potato amylopectin sample.

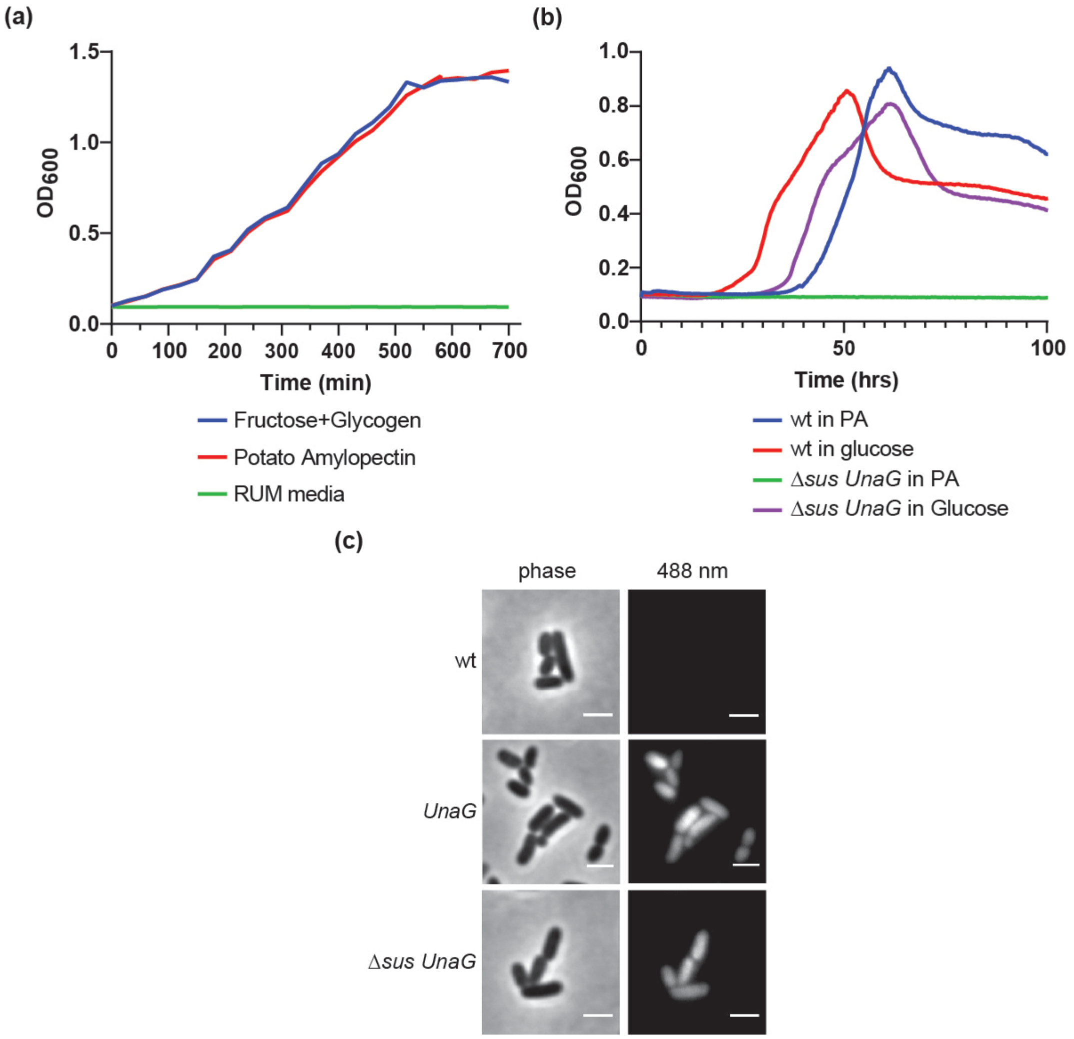


**Figure S3.** (a) Growth of *Rb* cells in RUM medium containing fructose (5 mg/ml) and glycogen (2.5 mg/ml) (blue), potato amylopectin (5 mg/ml) (red), and RUM medium control lacking added carbohydrate (green) measured by absorbance at 600 nm. (b) Growth of wt and Δ*sus UnaG Bt* cells in minimal medium containing potato amylopectin and glucose measured by absorbance at 600 nm. (c) Phase contrast and fluorescence imaging (488 nm channel) of wt, *UnaG*, and Δ*sus UnaG Bt* cells in the presence of bilirubin. All fluorescence images are normalized to the same brightness. Scale bars: 2 μm.

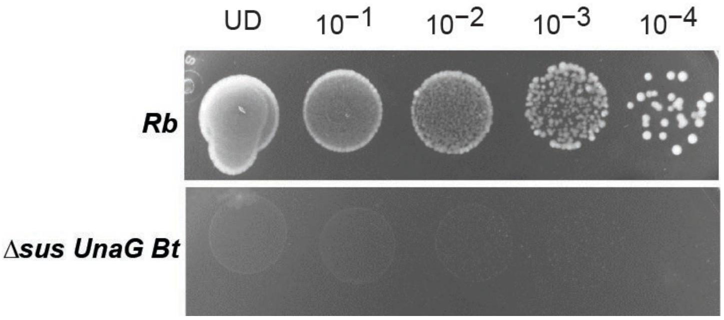


**Figure S4.** Pictures of *Rb* and Δ*sus UnaG Bt* cells plated on agarose plates at dilutions from undiluted (UD) to $10^{-4}$ with potato amylopectin (5 mg/ml). The same image contrast is used for both plates.

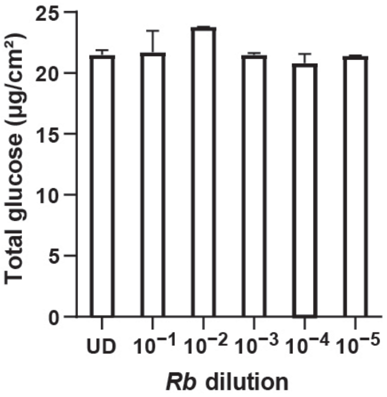


**Figure S5.** Total amount of glucose measured in the solid media taken from an area of 1 $cm^2$ within the halo around *Rb* across different dilutions (UD to $10^{-5}$). The average values of two biological replicates are shown. Error bars indicate standard error of mean.

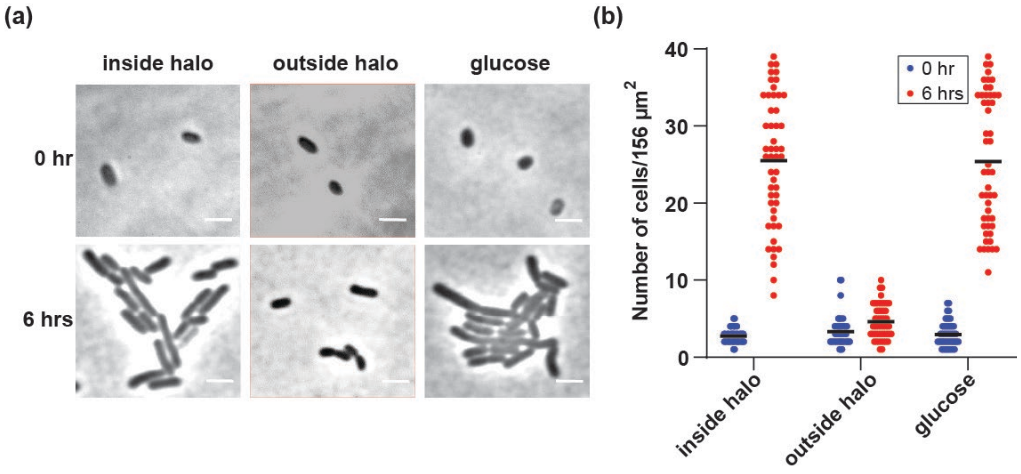


**Figure S6.** (a) Representative phase-contrast microscopy images of Δ*sus UnaG Bt* cells inside the glucose halo, outside the glucose halo, and on plates with glucose at 6 hrs. The 6 hrs time points from Figure 3a are repeated on the bottom panel for comparison. Scale bars: 2 µm. (b) Number of Δ*sus UnaG Bt* cells observed in each 12.5 µm × 12.5 µm imaging frame at 0 hrs (blue) and 6 hrs (red) inside the glucose halo, outside the glucose halo, and on plates with 125 µM glucose. Mean value is indicated by black line.

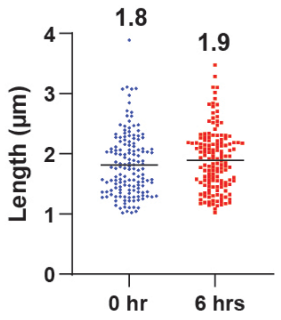


**Figure S7.** Length of Δ*sus UnaG Bt* cells grown in liquid medium containing glucose (27 mM) at 0 and 6 hrs. Each circle represents the length of a single cell from three biological replicates ($n$ = 150 cells). Mean value is indicated by black line. Statistical significance was determined by one-way ANOVA and post-hoc Tukey test. No statistically significant difference was seen between the samples.

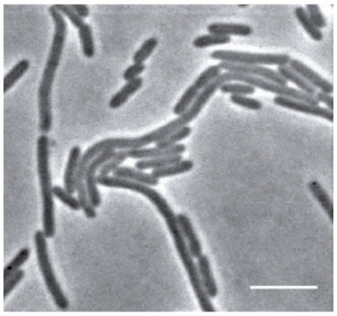

**Figure S8.** Representative phase-contrast microscopy image of wild-type *Bt* cells taken after 6 hrs incubation on agarose plates with 20 mM glucose. Scale bar: 5 µm.

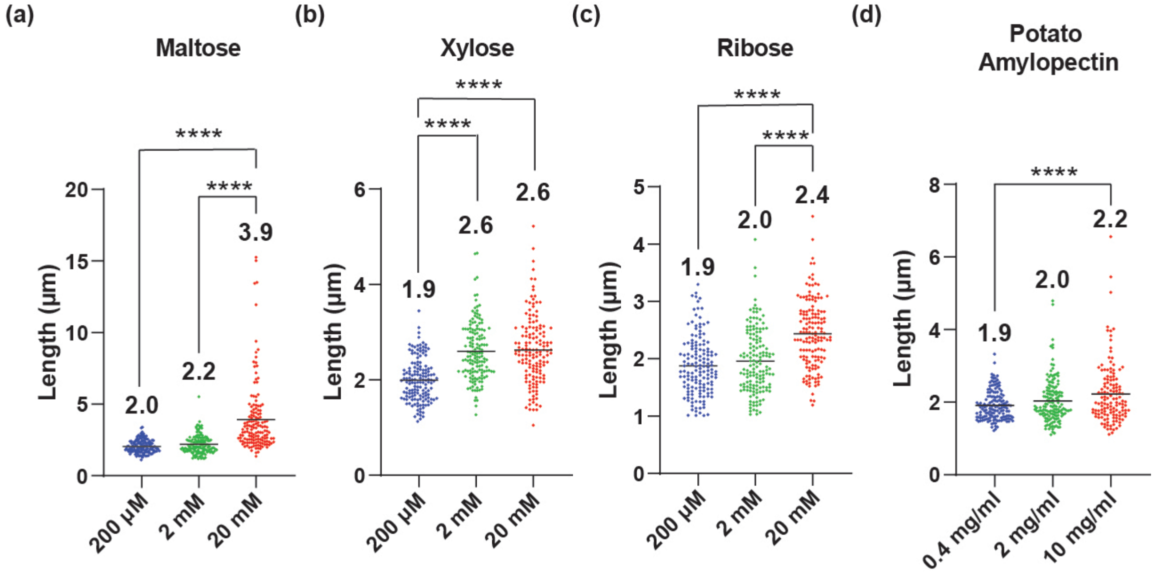


**Figure S9.** Length of wild-type *Bt* cells grown on 200 µM (blue), 2 mM (green) and 20 mM (red) of sugars after 6 hrs incubation on agarose plates with (a) maltose, (b) xylose, and (c) ribose. (d) Length of wild-type *Bt* cells grown on potato amylopectin at concentrations of 0.4, 2, and 10 mg/ml Each circle represents the length of a single cell from three biological replicates ($n$ = 150 cells). Mean value is indicated by black line. Statistical significance was determined by one-way ANOVA and post-hoc Tukey tests. Significant differences are indicated by **** ($P < 0.0001$).
